# *Buchnera* has changed flatmate but the repeated replacement of co-obligate symbionts is not associated with the ecological expansions of their aphid hosts

**DOI:** 10.1101/086223

**Authors:** A.S. Meseguer, A. Manzano-Marín, A. Coeur d’Acier, A-L. Clamens, M. Godefroid, E. Jousselin

## Abstract

Symbiotic associations with bacteria have facilitated important evolutionary transitions in insects and resulted in long-term obligate interactions. Recent evidence suggests that these associations are not always evolutionarily stable and that symbiont replacement and/or supplementation of an obligate symbiosis by an additional bacterium has occurred during the history of many insect groups. Yet, the factors favoring one symbiont over another in this evolutionary dynamic are not well understood; progress has been hindered by our incomplete understanding of the distribution of symbionts across phylogenetic and ecological contexts. While many aphids are engaged into an obligate symbiosis with a single Gammaproteobacterium, *Buchnera aphidicola*, in species of the Lachninae subfamily, this relationship has evolved into a *“ménage à trois”*, in which *Buchnera* is complemented by a cosymbiont, usually *Serratia symbiotica.* Using deep sequencing of 16S rRNA bacterial genes from 128 species of *Cinara* (the most diverse Lachninae genus), we reveal a highly dynamic dual symbiotic system in this aphid lineage. Most species host both *Serratia* and *Buchnera* but, in several clades, endosymbionts related to *Sodalis, Erwinia* or an unnamed member of the Enterobacteriaceae have replaced *Serratia*. Endosymbiont genome sequences from four aphid species*+*confirm that these coresident symbionts fulfill essential metabolic functions not ensured by *Buchnera.* We further demonstrate through comparative phylogenetic analyses that co-symbiont replacement is not associated with the adaptation of aphids to new ecological conditions. We propose that symbiont succession was driven by factors intrinsic to the phenomenon of endosymbiosis, such as rapid genome deterioration or competitive interactions between bacteria with similar metabolic capabilities.

## Introduction

Symbiotic associations with bacterial partners have facilitated important evolutionary transitions in the life histories of eukaryotes and have probably driven species diversification. Some groups of plant-eating insects have made use of the metabolic versatility of bacteria to feed on plant parts lacking certain essential nutrients (Hansen & Moran 2014). They generally shelter their bacterial partners within specialized cells and transmit them from mother to offspring (Buchner 1965). This type of nutritional endosymbiosis was first described in aphids (Hemiptera: Aphididae), a group of about 5000 species that feed on the phloem of their host plants. Almost all aphids host a Gammaproteobacterium, *Buchnera aphidicola*, which provides them with essential amino acids and vitamins that are rare in their diet (Douglas 1998; Wilson *et al.* 2010). Aphids may also be associated with facultative endosymbiotic bacteria that are not required for survival; their prevalence varies across populations (Oliver *et al.* 2010).

Obligate endosymbionts provide net benefits to their hosts, but reliance on long-term symbiotic associations can sometimes lead to evolutionary *“dead-ends”* (Bennett & Moran 2015). The maternal transfer of bacteria causes severe bottlenecks in bacterial populations, leading to genetic drift and the fixation of slightly deleterious mutations (Moran 1996; Rispe & Moran 2000; Toft & Andersson 2010). This process may alter symbiotic functions (McCutcheon & Moran 2012) and limit the thermal tolerance of bacteria (Wernegreen 2012), ultimately having a deleterious effect on the host dependent on these bacteria. One possible outcome of this situation is the replacement or the supplementation of the ancestral symbiont by a new one. Since the biosynthesis of amino acids are ubiquitous capabilities in bacteria, in some insect species, some of the facultative endosymbionts have become more than occasional partners, either entirely replacing the ancestral primary symbiont (Conord *et al.* 2008; Koga & Moran 2014; Smith *et al.* 2013; Toju *et al.* 2013), or persisting alongside it whilst taking on a subset of its functions (McCutcheon & Moran 2010; McCutcheon & von Dohlen; Takiya *et al.* 2006; Wu *et al.* 2006). An increasing body of evidence now shows that symbiont replacements have occurred repeatedly in insects, yet the factors favoring one symbiont over another in this evolutionary dynamic are still not well understood. It has been suggested that the acquisition of a new symbiont may not only provide the insect with a way of coping with the degradation of the genome of the primary symbiont, but may also confer new metabolic capabilities on the insect host (Koga & Moran 2014; Toenshoff *et al.* 2012). The acquisitions of these bacteria would then represent key innovations that allow their hosts to diversify in ecological niches that would otherwise be unavailable to them (Moran & Telang 1998). Studies on facultative symbionts in aphid populations, and insects in general, support this hypothesis; they generally increase the fitness of their hosts in specific environments and mediate ecological interactions (Frago *et al.* 2012; Henry *et al.* 2015; Oliver *et al.* 2010; Oliver & Martinez 2014; Russell & Moran 2006). However, because obligate nutritional symbioses remain more stable over evolutionary times than facultative ones, investigating the ecological factors that govern obligate symbiont turnover requires a wide taxonomic coverage and a solid phylogenetic framework.

Recent studies have reported the presence of labile di-symbiotic systems in aphids of the Lachninae subfamily. In *Cinara cedri, Cinara tujafilina* and *Tuberolachnus salignus, B. aphidicola* has lost the ability to synthesize the essential compounds riboflavin and biotin (in *C. cedri* and *T. salignus* it has also lost the ability to synthesize tryptophan), and these functions are now fulfilled by a former facultative endosymbiont, *Serratia symbiotica* (Gosalbes *et al.* 2008; Lamelas *et al.* 2011a; Lamelas *et al.* 2011b; Manzano-Marin & Latorre 2014; Manzano-Marin *et al.* 2016a). *Serratia* has thus become a co-obligate partner with a nutritional role complementary to that of *Buchnera* (Manzano-Marin *et al.* 2016a). It has been suggested that the riboflavin biosynthetic capability of *Buchnera* was lost in the ancestor of the Lachninae (Lamelas *et al.* 2011b; Manzano-Marin *et al.* 2016a). This implies that a coresident symbiont is now required by all members of the Lachninae for the system to survive. Characterizations of endosymbiotic bacteria in members of the subfamily including some *Cinara* spp. have shown that all the specimens studied so far harbor at least one additional bacterial endosymbiont alongside *Buchnera* (Burke *et al.* 2009; Jousselin *et al.* 2016; Lamelas *et al.* 2008)-while most species host *Serratia symbiotica*, some are associated with an alternative member of the Enterobacteriaceae. Observations of endosymbiont morphology and location in their hosts lend further support to the obligate aspect of the association with this new bacterial partner (Manzano-Marin *et al.* 2016b). Altogether these results also suggest that symbiont replacement has occurred in Lachninae. However, these studies were conducted on relatively few species and usually a single specimen per species. Results from a few samples represent mere snapshots of the ongoing evolutionary dynamics of these associations. A full understanding of the factors mediating symbionts replacements requires the analysis of the distribution of obligate symbionts across wide phylogenetic and ecological contexts. The aphid genus *Cinara* (Lachninae) might be an ideal model to conduct such a study. *Cinara* accounts for more than half of the Lachninae species diversity and is the second most diverse genus of aphids. It has diversified on various conifer genera, giving rise to more than 240 species (Chen *et al.* 2015; Meseguer *et al.* 2015). Species of this genus are distributed throughout the Holarctic and originated about 45 Mya, surviving all climatic changes that occurred through the Cenozoic (Zachos *et al.* 2008). Therefore, *Cinara* spp. have experienced a wide range of ecological conditions during their evolution. This long evolutionary history might have been accompanied by major changes in symbiotic interactions.

To elucidate the long-term evolution and maintenance of symbiotic associations in *Cinara*, we carried out an extensive survey of endosymbionts on a sample encompassing 50% of the genus’ known species diversity. We deep sequenced 16S rRNA genes, and modeled their distribution across the aphid phylogeny. We also sequenced the paired *Buchnera* and- *Serratia, Erwinia, Sodalis* or *Type-X* genomes (the main endosymbionts identified in our study) of four *Cinara* species to search for the presence/absence of the Riboflavin biosynthetic genes. We found that *Cinara* species have acquired different companion symbionts alongside *Buchnera* during the course of their diversification and that those have become obligate partners of the association complementing *Buchnera* in its nutritional role. We then explored the evolutionary pathways leading to the replacements of symbionts in this obligate dual symbiosis, by investigating whether the variation of host life-history traits and the climatic conditions experienced by the aphid were correlated with changes in co-obligate symbiont identity.

## Experimental Procedures

### 16S rDNA Endosymbiont characterization

#### DNA samples and 16S rDNA amplification

We sampled 366 colonies of *Cinara* and 5 outgroup colonies from the Lachninae and Mindarinae in the field. We sampled several colonies per species (from 1 to 17) to represent the species geographic distribution and diversity of host-plants. Aphids were kept in 70% ethanol at 6 °C immediately after collection. They were identified in the laboratory using different keys (Blackman & Eastop 2000; Favret & Voegtlin 2004). Collection details are given in Appendix 1. A single individual per colony was washed three times in ultrapure water and total genomic DNA was extracted from whole individuals with the DNeasy Blood & Tissue Kit (Qiagen, Germany), according to the manufacturer’s recommendations. The DNA was eluted in 40 sL of elution buffer. During the extraction procedure a negative control (i.e. a ‘blank template’ of ultrapure water) was processed with the same extraction kit. All DNA samples were stored at −20 °C. We amplified a 251bp portion of the V4 region of the 16S rRNA gene (Mizrahi-Man *et al.* 2013), and used targeted sequencing of indexed bacterial fragments on a MiSeq (Illumina) platform (Kozich *et al.* 2013), following the protocol described in (Jousselin *et al.* 2016). DNA extracts were amplified twice along with negative controls. PCR replicates were conducted on distinct 96-well microplates. As positive DNA controls, we used DNA extracts from three pure bacterial strains and three arthropod specimens with known bacterial endosymbionts.

We obtained a total of 749 PCR products, which were pooled and submitted for paired-end sequencing on a MISEQ (Illumina) FLOWCELL equipped with a version 2, 500-cycle reagent cartridge.

#### Sequence analyses and Taxonomic assignation

We used Mothur v1.3.3 (Schloss & Westcott 2011) implemented on a Galaxy workbench (Goecks *et al.* 2010) (http://galaxy-workbench.toulouse.inra.fr/) to assemble paired-end reads and filter out sequencing errors and chimeras from the results. The overlapped paired-end reads were assembled with the *make.contigs* function of MOTHUR, and the contigs exceeding 280 bp in length and/or containing ambiguous base pairs were filtered out and excluded from further analyses, since the V4 region is expected to have about 251 pb. A FASTA file containing unique contigs and a file reporting the occurrence of these sequences in each sample were created. Unique sequences from the FASTA file were then aligned with the V4 portion of reference sequences from the SILVA 16S reference database (v119) (Quast *et al.* 2013). Sequences that did not align with the V4 fragment were excluded from further analyses. After this filtering step, a new file containing unique sequences was created. The number of reads resulting from sequencing errors was then reduced by merging rare unique sequences with frequent unique sequences with a mismatch of no more than 2 bp relative to the rare sequences *(pre.cluster* command in MOTHUR). We then used the UCHIME program (Edgar *et al.* 2011) implemented in MOTHUR to detect chimeric sequences and excluded them from the data set. For each sequence, the number of reads per sample was transformed into percentages using an R script (Jousselin *et al.* 2016) and used to compile a contingency table (Appendix 2). We removed individual sequences representing less than 0.5 % of the reads in each sample. Sequences represented by such a small proportion of the reads were often found in negative controls, were generally not arthropod endosymbionts and, in most cases, were not found across PCR replicates of the same sample, suggesting that they could represent contaminants or spurious sequences (Jousselin *et al.* 2016). For each sample, we then eliminated all the sequences that did not appear in both PCR replicates.

Taxonomic affiliations of each unique sequence were obtained using the RDP Classifier in Qiime (Caporaso *et al.* 2010), with the Silva database, and leBIBI^QBPP^ (Flandrois *et al.* 2014), with the 16S SSU-rRNA-TS-stringent database. In addition, a neighbour-joining tree was reconstructed with all unique sequences combined with sequences of aphid endosymbionts identified in previous studies; sequences of *Hamiltonella, Phlomobacter, Erwinia, Dickeya, Edwardsiella, Sodalis, Pantoea, Klebsiella, Spiroplasma* and *Cardinium* were retrieved from Silva. Sequences of *Regiella* identified in a previous study (Smith *et al.* 2015), and sequences of *Arsenophonus, Buchnera, Hamiltonella*, the secondary symbionts of *Cinara* spp. identified by (Burke *et al.* 2009) were retrieved from NCBI. We then checked the coherence of the phylogenetic clusters obtained with the NJ tree and the taxonomic assignation from RDP and leBIBI.

#### Assessing endosymbiont diversity and specificity across samples

We added the frequencies of unique sequences assigned to a particular bacterial genus (or higher rank when genus assignation was not available) in a sample. We assessed the replicability of our results by plotting the percentage of reads assigned to each bacterium in one PCR replicate against the other for all *Cinara* samples, and calculated the Pearson correlation coefficient. We then plotted the bacterial community (phylum and frequency as estimated by read abundances) of each *Cinara* specimen/species to the tips of the *Cinara’s* phylogenetic tree (see below) using the R package *gplots v2.23* (Warnes *et al.* 2015); for the species-level analyses, we only considered the bacteria that were found in all the specimens of a given species. We calculated the mean prevalence of each endosymbiont in each *Cinara* species as the percentage of specimens per species that harboured a particular symbiont against the total number of individuals collected for that aphid species, excluding outgroups and species that were represented by a single specimen to avoid overestimation of mean values.

We represented the specificity of the aphid-endosymbiont associations by plotting the links between 16S bacterial sequences and the *Cinara* specimens/species in which they were found using *bipartite* (Dormann *et al.* 2008).

### Symbiont genome

#### Endosymbiont DNA extraction and sequencing

In order to obtain genome data of putative co-obligate endosymbionts of *Cinara*, for four species *(Cinara confinis, C. pseudotaxifoliae, C. strobi, C. fornacula)*, we prepared DNA samples enriched with bacteria following a slightly modified version of the protocol by Charles and Ishikawa (Charles & Ishikawa 1999) as described in Jousselin et al (2016). For this filtration procedure, for each aphid colony, 7 to 15 aphids were pooled together. DNA libraries were then prepared using the Nextera XT Library Kit (Illumina) and each library was multiplexed and sequenced as a combination of 300bp paired-end and/or 250 paired-end read on MiSeq (Illumina) flowcells and/or 100bp paired end read on one fourth of an Illumina Hiseq2000 lane (see supplementary information for details on the sequencing efforts used for each sample).

#### Draft genome assembly and annotation of riboflavin biosynthetic genes

Before assembly, reads were quality-trimmed using FASTX-toolkit v0.0.14 (http://hannonlab.cshl.edu/fastx_toolkit/) and reads shorter than 75 bps were filtered out. Additionally, reads containing undefined nucleotides (“N”) were discarded using PRINSEQ-lite v2.24.0 (Schmieder & Edwards 2011). Remaining paired reads were used for *de novo* assembly in SPAdes v3.8.0 (Bankevich *et al.* 2012) with kmer lengths of 33, 55, 77 for samples “2801” and 33, 55, 77, 99, 127 for samples “3056” and “3249”. The resulting contigs were then taxonomically assigned to a putative symbiont through blastx searches against a database composed of the proteome of the pea aphid (ABLF00000000), diverse aphid endosymbiont bacterial strains *(Buchnera* spp. [BA000003, AP001070-1, AE013218, AE016826, AF492591, CP000263, AY438025, EU660486] *Hamiltonella defensa* [CP001277-8], *Serratia symbiotica* [CP002295, CCES00000000, FR904230-48, HG934887-9], *Regiella insecticola* [ACYF00000000]), *Sodalis* spp. (CP006569-70, AP008232-5, CP006568), *Wolbachia* spp. (AM9998877, AP013028), and *Erwinia* spp. (FN666575-7, FP236842, FP236827-9, FP928999) strains, followed by manual curation. Afterwards, the resulting references were used for mapping with bowtie v2.2.5 (Langmead & Salzberg 2012) and reassembled using SPAdes (as previously described). Draft *Buchnera* chromosome assemblies were scaffolded using *Buchnera* from *Cinara tujafilina* (GenBank:CP001817.1) as reference. Final contigs were manually curated to remove spurious sequences (resulting from misassembles or contamination). Riboflavin biosynthetic genes were searched for using the online tblastn server, followed by manual curation.

### Phylogenetic relationships in *Cinara*

#### DNA sequences and phylogenetic reconstruction

We used five DNA fragments to reconstruct the phylogeny of the 371 aphids used in the endosymbiont survey. We used three DNA fragments from the aphid’s genome (cytochrome c oxidase subunit I *“COI”;* cytochrome b *“Cytb”;* and the elongation factor “EU”) and two from the DNA of *Buchnera aphidicola* (a chaperonin assisting in the folding of proteins *“GroEL”;* and *“His”*, which includes the ATP phosphoribosyltransferase (HisG) gene, the histidinol dehydrogenase (HisD) gene and the intergenic region). Total genomic DNA was extracted from a single individual from the same aphid colony as used in the endosymbiont survey. DNA was extracted, amplified and sequenced as in previous studies (Jousselin *et al.* 2013). A total of seventy-five specimens were newly sequenced for this study, while the other sequences were retrieved from GenBank (Appendix 1). Contigs were assembled from forward and reverse reads and corrected with Geneious 8.1.7 (Drummond *et al.* 2010). Alignments were generated with *mafft* v.6 (Katoh & Toh 2008), with the default option L-INS-I, and were manually adjusted with *Se-Al* 2.0a11 Carbon (Rambaut 2002).

We concatenated all the markers in a single matrix and inferred phylogenetic relationships using MrBayes v3.2 (Ronquist *et al.* 2012) in a dataset including the sequences of 371 aphid individuals *(specimens dataset)* and a reduced dataset *(species dataset)* including only one specimen per phylogenetic cluster retrieved in the delineation test (128 species; see below). We evaluated different partitioning strategies of the datasets using Bayes factor comparisons of the harmonic mean to determine the best-fit partitioned scheme: *UnPart* (single data partition), *GenePart* (partitioned by gene), *PartFind* (partitioned following Partition Finder results), and *MixedPart* (the mixed model described below). For the *PartFind* scheme, we used the partitioning scheme and across-site rate variation suggested by PartitionFinder v1.1 (Lanfear *et al.* 2012); we inferred the best substitution model for each partition among those available in MrBayes, using the Bayesian Information Criterion (BIC) metric under a *greedy* algorithm. For the *MixedPart* scheme, instead of *a priori* applying a specific substitution model for each partition, we sampled across the substitution model space using a reversible-jump Markov Chain Monte Carlo (rj-MCMC), with the option *nst=mixed.* This procedure integrates the uncertainty concerning the correct structure of the substitution model (Huelsenbeck *et al.* 2004). The best-fit partitioned scheme was the *PartFind* (supplementary information Table S1), which was used for subsequent analyses. To prevent the overestimation of branch lengths when mutation rates differ between partitions of different genes (Brown *et al.* 2010) and between regions of single genes (Brown *et al.* 2010; Meseguer *et al.* 2013), we set the value of the shape parameter λ, which controls the exponential prior for branch lengths, to *λ* =100, assigning greater probability to short branches. We conducted 2 independent runs of 4 Metropolis-coupled chains each for 40 million generations, sampling every 1000 generations and discarding 20% as burnin.

#### Aphid species delimitation

Aphids show considerable overlap in their morphological characters; consequently, their identification often relies on biological traits such as host-plant associations, which renders further investigations on the evolution of aphid life-history traits tautological (Coeur d’acier *et al.* 2014). To avoid this bias, we complemented the morphological identifications of specimens with DNA-based species delimitation analyses. We used the Bayesian implementation of the Poisson tree processes (BPTP) model (Zhang *et al.* 2013) to delimit putative *Cinara* species. We ran the analysis for 500.000 generations, thinning every 100, and discarding 0.1 % as burnin. We considered the clusters retrieved in this analysis as “phylogenetic species” and repeated the phylogenetic analyses including only one individual per phylogenetic species.

### Phylogenetic comparative analyses

We tested whether endosymbiotic associations were phylogenetically conserved using the *λ* of *Pagel* (Pagel 1994). It is a quantitative measure that varies between 0 (when there is no phylogenetic signal in the trait) and 1 (when there is phylogenetic signal). We optimized the value of lambda for the presence/absence of each bacterial lineage found in our samples onto the species tree using maximum-likelihood (ML) in *geiger* (Harmon *et al.* 2008). The presence/absence of *Serratia, Erwinia, Sodalis, Wolbachia, Hamiltonella, Type-X*, and *Acetobacteraceae* were modelled as binary traits. We did not analyse other endosymbionts detected in our study since they were poorly represented in our samples nor fixed within *Cinara* species (i.e. they never infected all the individuals of the same species). The species-level analysis excluded *Rickettsia, Regiella* and *Arsenophonus* that were fixed in few species. To test if *λ* was significantly different from 0 we compared a model with the observed value of *λ* to a model with a fixed *λ* of zero using a likelihood ratio test.

We inferred ancestral associations of *Cinara* with each bacteria showing phylogenetic signal using ML in *ape* (Paradis *et al.* 2004) over the species phylogeny. Symbiotic associations were treated as a discrete character with 7 states: *Serratia, Erwinia, Sodalis, Type-X, Wolbachia, Hamiltonella* and “No cosymbiont”, for species in which no bacterium, apart from *Buchnera*, was fixed in the species. Transition probabilities between character states were estimated under two models; equal rates “ER” and all-rates different “ARD” model, where all possible transitions between states receive distinct parameters. A likelihood ratio test was used to select the most appropriate model.

We evaluated the factors that were correlated with the presence of symbionts across *Cinara* species using logistic phylogenetic regressions (Ives & Garland 2010) in *phylolm* (Ho & Ané 2014). We fitted five different regression models for each of the bacteria exhibiting phylogenetic signal, with each time its presence/absence as the dependent variable and one explanatory variable. We assessed the significance of the correlations by comparing these models with a null model using AIC. We chose explanatory variables that reflect several dimensions of the aphid’s ecological niches and are generally used to explain the distribution of secondary symbionts across aphid populations. We included three life-history traits related to host-plant use: *(i) host-plant genera;* i.e. *Pinus, Picea*, Cupressaceae, *Larix, Pseudotsuga, Abies* or *Cedrus, (ii) feeding range;* whether species were monophagous (feeding on a single plant species or a few closely related species) or polyphagous, and *(iii) feeding site*; whether species fed on lignified parts of the plant (branches and trunks) or not (needles, shoots, young twigs or at the base of new cones). These characters likely reflect the panel of variations in the metabolic needs of aphid species. We also tested *(iv)* the role of *aphids’ life habit;* i.e. whether aphids lived solitarily or in dense colonies. Differences in life habit as well as variations in host-plant use are likely associated with variations in the communities of natural enemies of aphids, which might favour alternative defensive symbionts (Cayetano & Vorburger 2015; Henry *et al.* 2015;

Oliver *et al.* 2008; Smith *et al.* 2015). We assigned character states by combining information available in the literature for each recognized *Cinara* species (Blackman & Eastop 1994; Jousselin *et al.* 2013) and information recorded from the field in the course of aphid sampling. We did not explore the effect of aphid life cycle nor the association of aphids with ants as all *Cinara* species are monoecious and almost all are attended by ants. Recent studies underlined that expansion to new geographic areas could favour the acquisition of new bacterial partners in aphids (Zytynska & Weisser 2016), we therefore investigated whether the *(v) aphids’ geographic distribution-* i.e. whether species were distributed in the western or eastern parts of the Nearctic and the Palearctic-could explain variations in symbiont partnerships. Widespread geographic ranges in species of *Cinara* mostly resulted from recent dispersal events; we thus coded the distribution of these species according to the distribution of their most recent ancestor estimated in a previous study (Meseguer *et al.* 2015). The prevalence, distribution and abundance of symbionts (both obligate and facultative) across aphids can also vary with the temperature (Russell & Moran 2006), suggesting that symbiont turnover could be driven by climatic variations. We thus tested the effect of climatic variables in the distribution of symbionts. Climatic values of specimen records were extracted from 6 raster layers, Worldclim (Hijmans *et al.* 2005), at a resolution of 30 arc-seconds: annual mean temperature, temperature of the coldest and the warmest month, annual precipitation, and precipitation of the wettest and driest month. Multiple occurrences of a symbiont in the same grid cell were reduced to a single occurrence. We ran between-group principal component analysis (PCA) (Dolédec & Chessel 1987) to compare climatic envelopes of symbionts on *ade4* (Dray & Dufour 2007). We tested the significance of the between-groups structure using Monte-Carlo permutation tests with 999 replications. Temperature and precipitation ranges of symbionts were also visualized with barplots.

## Results

### 16S rDNA dataset description

High-throughput sequencing of 16S rRNA bacterial genes from 371 individual aphids generated 8.412.145 reads passing Illumina stringent quality control (mean number of reads per sample=11.246, standard deviation=9113; excluding negative controls). After all sequence filtering steps, we obtained 8.403.870 reads, corresponding to 182.913 unique sequences (Appendix 3). After discarding sequences accounting for less than 0.5% of the reads in each sample, we obtained 630 unique sequences. Overall, 88.7% of these sequences were attributed to Enterobacteriaceae *(Arsenophonus, Buchnera, Edwardsiella, Erwinia, Hamiltonella, Regiella, Serratia, Sodalis)*, 2.7% to Rickettsiaceae *(Rickettsia, Wolbachia)*, 1.6% to Acetobacteraceae, 0.6% to *Acinetobacter* and 0.3% to *Spiroplasma.* The remaining 6% of the sequences were assigned to families containing water-and soil-borne bacteria *(e.g.* Comamonadaceae, Flavobacteriaceae or Methylobacteriaceae), each of which occurred at very low frequency (Appendix 2). The removal of sequences accounting for less than 0.5% of the reads in aphid samples eliminated most of the sequences common to negative controls (Fig. S1). The bacterial taxonomic compositions of aphid samples were highly similar across PCR replicates (r^2^>0.99; Fig. S2), except for *Acinetobacter* (r^2^<0.8) which did not appear in similar proportions in replicates.

### Diversity of symbionts associated with Cinara

Of the 371 aphid specimens examined here, 218 hosted two bacteria: *B. aphidicola* and a second partner belonging to various lineages: *Serratia, Erwinia, Sodalis, Wolbachia* or a non-described lineage of Enterobacteriaceae. The 16S rDNA sequence of the latter is highly similar to the one of the secondary symbiont found associated with *Acyrtosiphon pisum*, named *Type-X* (Guay *et al.* 2009). We thus referred to this symbiont as *Type-X* throughout the manuscript (Fig. 1, S3; Table S2). 146 specimens hosted *Buchnera* alongside with 3-5 other bacterial partners). *Serratia, Erwinia, Sodalis, Wolbachia* and *Type-X* tended to be present in all the individuals of the species they infected, with an overall mean prevalence for each bacterium over 70% (i.e. each bacterium appeared, on average, in more than 70% of the specimens studied) (Fig. S4). Conversely, *Arsenophonus, Hamiltonella, Regiella*, *Rickettsia, Acinetobacter* and Acetobacteraceae were generally detected in some, but not all of the individuals of the species they infected (the overall mean prevalence for each of these bacteria was below 60%). Seven *Cinara* specimens did not contain an alternative bacteria along with *Buchnera*: *C. laricis* (1 specimen out of the 3 studied was associated only with *Buchnera), C. kochi* (1/1), *C. brevispinosa* (2/11), *C. cfcoloradensis* (1/2), *C. close piceae* (1/4) *and C. sp_3210* (1/1).

**Figure 1.**
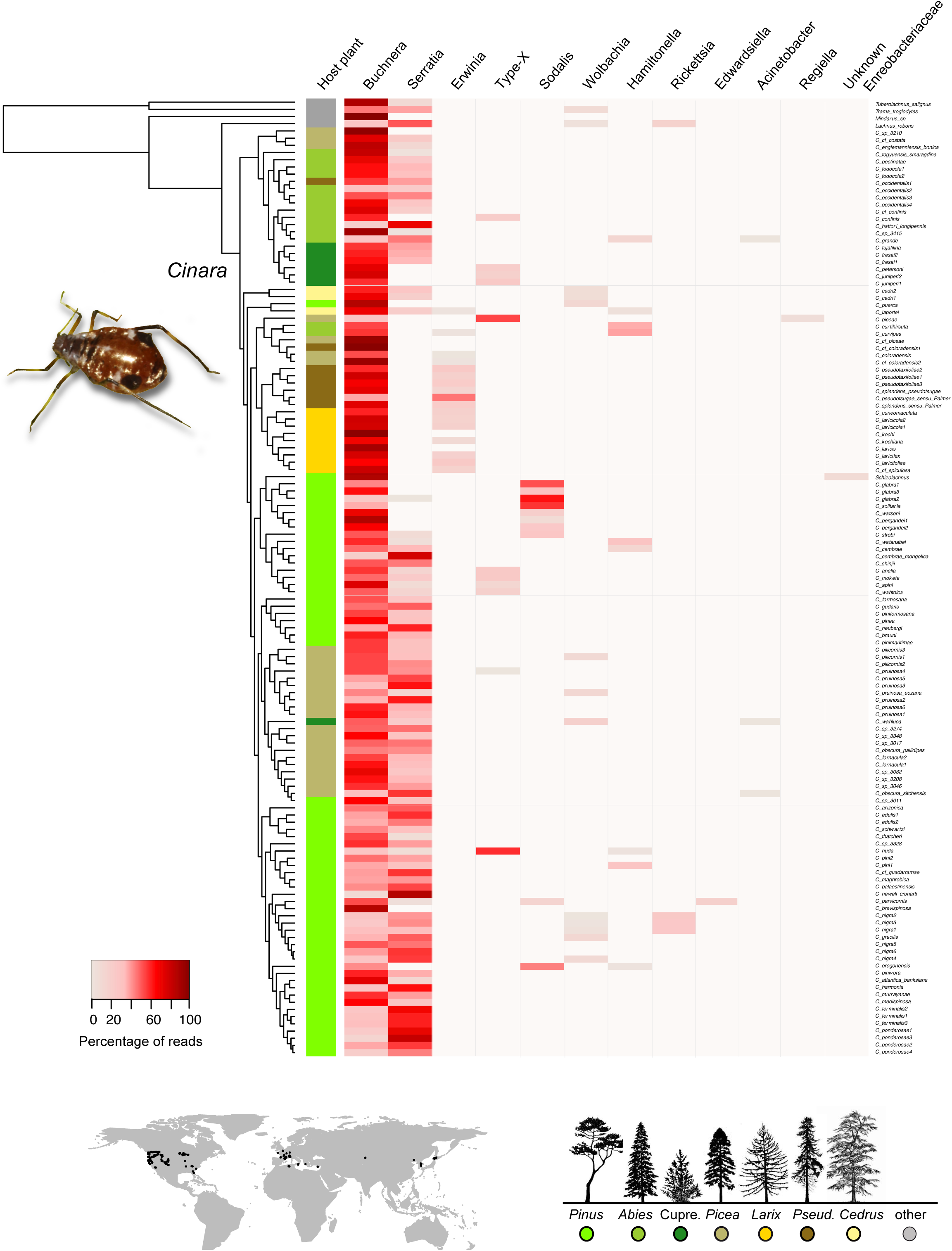
Bacterial community (phylum and frequency as estimated by read abundances) of *Cinara* species mapped onto the tips of the ultrametric species tree of *Cinara*. For the total number of reads found in an aphid specimen, we have calculated the percentage of reads belonging to each bacterium. In this figure, only endosymbionts that are fixed within *Cinara* species (bacteria found in all specimens of a given species) are plotted. Therefore, the frequency of each endosymbiont (the red bars) in a species represents the mean percentage of reads obtained in all the specimens of that species (including PCR replicates). Colours in the figure correspond to the colour circles and conifer silhouettes in the inset legend. Black circles in the inset map show the collection sites of the specimens of *Cinara* (N=371). The photo shows *Cinara ponderosae* (col: Coeur d’acier).Colours in the figure corre-the colour circles and conifer silhouettes in the inset legend. Black circles in the inset map show the collection sites of the is of *Cinara* (N=371). The photo shows *Cinara ponderosae* (col: Coeur d’acier).

### Symbiont specificity

The association between *Serratia* and *Cinara* was generally species-specific, with 68% of *Cinara* species hosting a single *Serratia* 16S rRNA gene sequence, and 74% of *Serratia* 16S rRNA gene sequence found associated with a single *Cinara* species (Fig. S5). These patterns of species specificity were very similar to those observed for *Cinara* and *Buchnera* (Table 1; Fig. S6). For the species in which the presence of more than one *Serratia* haplotype was reported, some cases resulted from different bacterial haplotypes being present in individuals from different populations, whereas in other cases, up to two *Serratia* haplotypes were found in a single *Cinara* specimen (Appendix 2). These situations may represent cases of co-infection of an aphid with different *Serratia* strains or they might stem from slight divergences in 16S rDNA sequences from a single *Serratia* strain (slightly divergent bacterial chromosomes can sometimes occur within a single bacteriocyte (Komaki & Ishikawa 1999). More than half of the *Cinara* species infected with Sodalis-related bacteria or *Type-X* were found associated with more than one haplotype of these endosymbionts (Table 1; Fig. S5), a single specimen could host up to 7 different 16s rDNA sequences (Appendix 2) (again indicating coinfections or multiple 16S rDNA sequences in a single bacterial strain) whereas about 90% of these bacteria were specific to their host (i.e. found associated with a single *Cinara* species). Overall, 85% of the *Cinara* species infected with *Erwinia* contained a single haplotype, and 50% of the *Erwinia* haplotypes were host-specific (Fig. S5). For all these bacteria, there were cases in which the same haplotype was found to be present in different *Cinara* species. Shared haplotypes were generally found in closely related *Cinara* species, but a few haplotypes were well represented in distantly related species (green lines in Fig. S5). We identified one *Serratia* 16S rRNA gene sequence that was present in 34 distantly related species of *Cinara.*

**Table 1.**
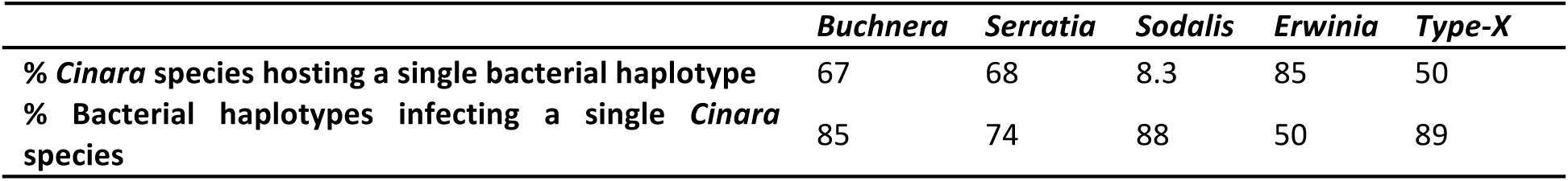
Aphid-symbiont specificity. Only well represented haplotypes (representing >1% of the reads in a sample) are considered.

| *Buchnera* | *Serratia* | *Sodalis* | *Erwinia* | Type-X |
| --- | --- | --- | --- | --- |

|  |  |  |  |  |  |
| --- | --- | --- | --- | --- | --- |
| % *Cinara* species hosting a single bacterial haplotype | 67 | 68 | 8.3 | 85 | 50 |
| % Bacterial haplotypes infecting a single *Cinara* species | 85 | 74 | 88 | 50 | 89 |

### Endosymbiont genomes

In order to corroborate the role of *Buchnera’s* co-resident symbionts as the riboflavin providers, we sequenced the genomes of *Buchnera* and the secondary symbiont of *C. strobi (Sodalis-like), C. fornacula (S. symbiotica), C. pseudotaxifoliae (Erwinia)*, and *C. confinis (Type-X).* The genomes of all *Buchnera* symbionts were assembled into one single circular scaffold plus the typical *leucine* and *tryptophan* plasmid.T genome of the *Erwinia-like* symbiont was assembled into one contig plus one circular plasmid and the genome of the *Serratia* was assembled into 1 contig. The genomes of the *Sodalis-like* and *Type-X* secondary symbionts were highly fragmented, given the high presence of mobile genetic elements. All four secondary symbionts hold small genomes, when compared to their free-living relatives, with the genomes of both *Erwinia (circa* 1.09Mb) and *Serratia (ca.* 1.16Mb) symbionts being the smallest. Full statistics for the genome assemblies can be found in Table S3. Blastx searches for *riboflavin, tryptophan* and *biotin* biosynthetic genes revealed that none of the *Buchnera* strains is able to synthesize riboflavin while all co-resident symbionts preserve intact routes for the biosynthesis of this compound (Fig. 2).

**Figure 2.**
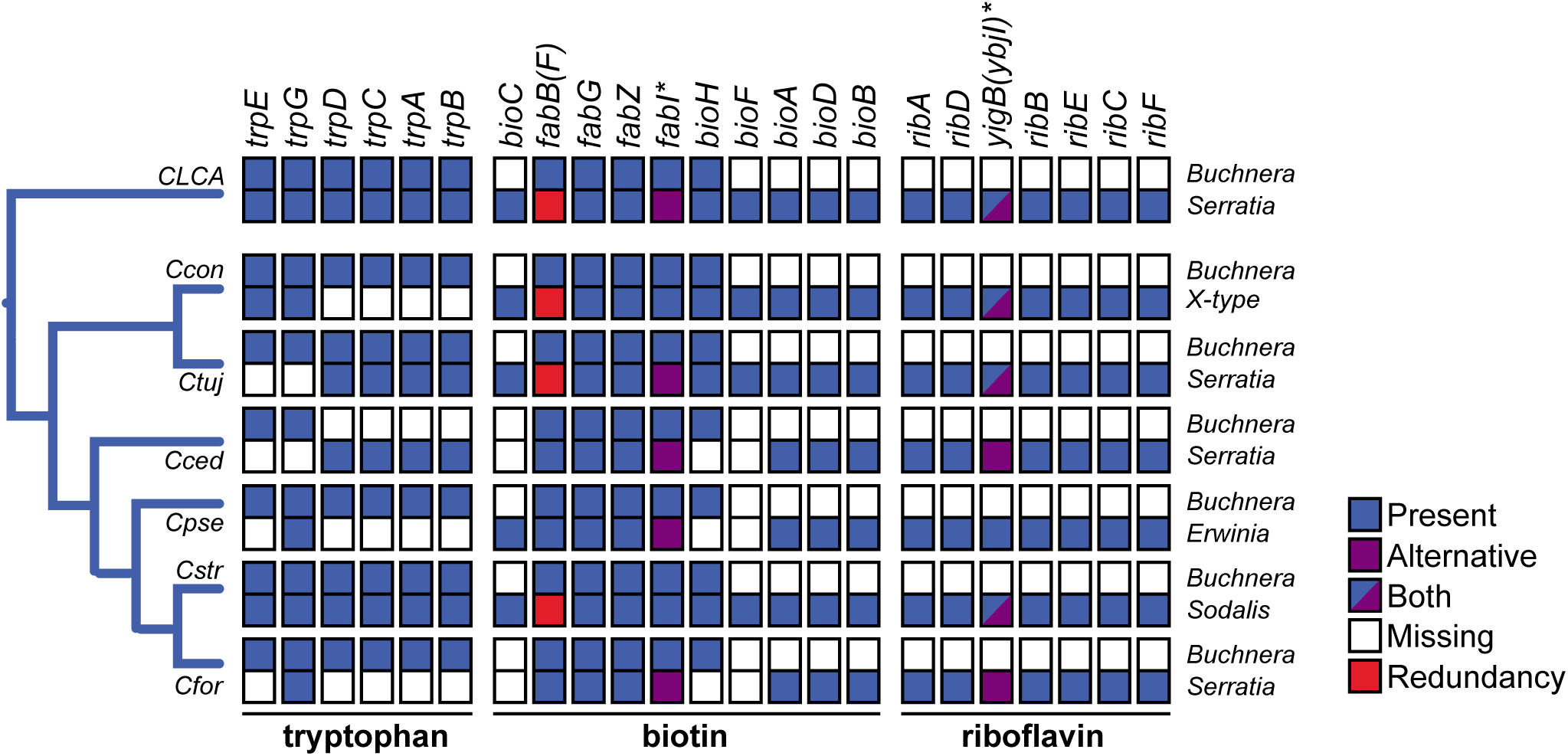
Diagram representing the metabolic complementation found in the biosynthesis of tryptophan, biotin and the riboflavin biosynthesis take-over/rescue between *Buchnera* and- *Serratia, Erwinia, Sodalis or Type-X* obligate symbionts in the endosymbiotic systems of *Cinara.* Gene names are used as column names. On the left, a schematic representation of the phylogenetic relationships of *Cinara* species. Abbreviations for the different species are: CLCA= *Cinara* last common ancestors; Ccon = *Cinara confinis*, Ctuj = *C. tujhafilina*; Cced = *C. cedri;* Cpse = *C. pseudotaxifoliae;* Cfor = *C. fornacula.* Data from C. *tujhafilina* and *C. cedri* comes from Gosalbes et al (2008), Lamelas et al. (2011a,b) and Manzano-Marín & Latorre (2014).

### Phylogenetic reconstruction of Cinara and species delimitation

The phylogeny of 371 aphid individuals was well resolved and highly sustained (pp >95) (Fig. S7). Species delimitation analyses with BPTP identified 128 putative *Cinara* species (acceptance rate=0.18; range: 121-147) among the total of 371 specimens analyzed, separating morphological species into several clusters in a few cases (Fig. S7). The phylogeny including one specimen of each of the 128 species retrieved in the delineation test was also well resolved and strongly supported (Fig.S8).

### Comparative phylogenetic methods

*Pagel’s λ* test indicated a significant (p<0.01) phylogenetic signal over the species tree for the association with *Serratia, Erwinia, Type-X, Wolbachia* and *Sodalis* (Table S4). The ancestral character state reconstructions suggested that *Serratia* was acquired in the common ancestor of all Lachninae (Fig. S9). *Serratia* was then independently lost in eight lineages, and recently reacquired in *Cinara glabra. Sodalis, Wolbachia, Erwinia* and *Type-X* were acquired more recently in different lineages of *Cinara*, following the loss of *Serratia*, although *Type-X and Wolbachia* persisted alongside *Serratia* in a small group of species.

The presence of different cosymbionts across *Cinara* species could not be explained by any of the aphid traits examined here, as the null models always fitted the data better than the models including explanatory variables (Table S5)-all of the obligate cosymbionts identified here were found in aphid species using a wide range of ecological niches (Fig. 3). The climatic envelopes of the main co-symbionts were not significantly different (inertia: 0.009; *P* = 0.298; Fig. 4, S10)

**Figure 3.**
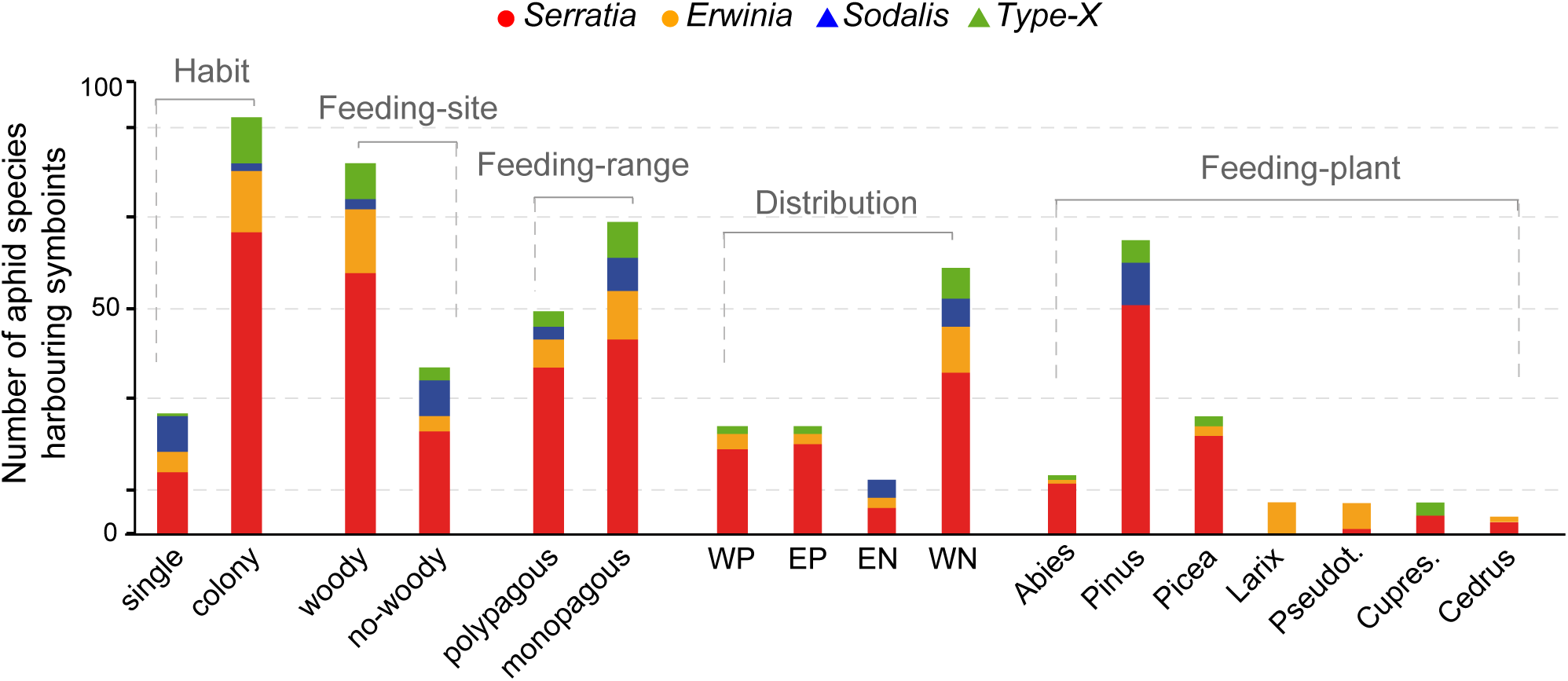
Distribution of cosymbionts across aphid’s ecological contexts. For several aphids characters, the histograms show the number of species hosting each cosymbionts. Abbreviations: Pseudot.= Pseudotsuga, Cupre.=Cupressaceae, WP= Western Palearctic, EP= Eastern Palearctic, WN= Western Nearctic, EN= Eastern Nearctic.

## Discussion

### Repeated evolution of dual symbiosis in Cinara

The evolutionary history of *Cinara* has been accompanied by major changes in its symbiotic partners. We show that, during the course of its diversification, *Cinara* has acquired different cosymbionts that reside with their primary symbiont, *Buchnera aphidicola.* In 79% of the 128 species studied here, *Buchnera* was found to coexist with *Serratia*, whereas, in the remaining 21%, *Serratia* was replaced by a bacterium related to *Erwinia, Sodalis* or an unknown member of the Enterobacteriaceae referred to here as *Type-X* (Fig. 1, Table S2). The tendency of these bacteria to be present in all the specimens of the species they infect (Fig. S4), throughout their geographic range and on their various host-plants, together with the phylogenetic conservatism of the associations (Table S4), suggests that they have established long-term relationships with their aphid hosts. The analyses of the endosymbiont genomes confirm that none of the *Buchnera* strains newly sequenced here is able to synthesize *riboflavin*, while all secondary symbionts preserve intact routes for the biosynthesis of this compound and would be capable of putatively complementing the truncated *biotin* biosynthetic pathway (Fig. 2). Altogether, these results, along with recent studies within Lachninae (Manzano-Marin *et al.* 2016b) add to the mounting evidence that all *Cinara* species and probably all Lachninae shelter a di-symbiotic system: *Serratia, Erwinia, Sodalis and Type-X* coexist with *Buchnera* as co-obligate partners. The absence of *riboflavin* biosynthetic genes in all currently sequenced *Buchnera* from *Cinara* aphids (Lamelas *et al.* 2011b; Manzano-Marin *et al.* 2016a; Perez-Brocal *et al.* 2006) suggests that the *Buchnera* from the *Cinara* last common ancestor (CLCA) was probably dependent on a secondary endosymbiont for the biosynthesis of this essential vitamin. Also, given the *Buchnera* gene content, it is highly likely that the CLCA developed a *biotin* auxotrophy: the biosynthesis of this vitamin could be split between *Buchnera* and its companion symbiont (Fig. 2). The finding that *Buchnera* seems to lack a co-resident symbiont in a few specimens (7 individuals in our sampling, Fig. S3) could challenge this interpretation-it could suggest that some individuals rely only on *Buchnera* and thus that the primary symbiont genome is still fully functional or that the aphid can fulfill the functions lost in *Buchnera;* it could have acquired this ability through horizontal gene transfer. Alternatively, the failure to detect a co-symbiont in these individuals could be the result of bacteriocyte size reduction and/or symbiont degradation that occurs with aphid aging. This phenomenon has been described for *Buchnera* bacteriocytes in the pea aphid (Simonet *et al.* 2016). It is also possible that individuals lose their co-obligate symbiont during their development; recent studies have shown that the cereal weevil *Sitophilus pierantonius*, eliminates its obligate symbiont (*Sodalis pierantonius*) when it no longer needs it (Vigneron *et al.* 2014). A thorough investigation of symbiont cell localization within their host and dynamics throughout the aphid development will be needed to validate any of these interpretations. Specimens without a co-symbiont mostly belonged to the *Cinara* clade associated with *Erwinia;* it will be interesting to follow bacteria cell population dynamics of this symbiotic association.

The obligate association of *Serratia* with *C. tujafilina, C. cedri* and *Tuberolachnus salignus* has already been demonstrated through genome-based metabolic inference (Lamelas *et al.* 2011b; Manzano-Marin & Latorre 2014; Manzano-Marin *et al.* 2016a). Associations of aphids and bacteria related to *Sodalis* have rarely been documented. However specific PCR assays and histological work in Lachninae (Burke *et al.* 2009; Manzano-Marin *et al.* 2016b) have shown that Sodalis-related bacteria were associated with several species, and suggested that these bacteria could have replaced *Serratia* in different Lachninae lineages. Sodalis-related bacteria are actually ubiquitous in insects and have seemingly established obligate associations with various hemipterans (Husnik & McCutcheon 2016; Koga *et al.* 2013; Koga & Moran 2014; Oakeson *et al.* 2014; Snyder *et al.* 2011). *Type-X* has also been detected in members of the Lachninae and its presence was interpreted as a replacement for *Serratia* in some lineages (Manzano-Marin *et al.* 2016b). Bacteria related to *Erwinia* are generally free-living plant pathogens. However, it has been suggested that they can act as symbiotic partners of aphids (Clark *et al.* 2012; Harada *et al.* 1997), though it is usually assumed to be an arthropod gut symbiont. This is the first study to show an *Erwinia* related lineage to be an obligate symbiotic partner of aphids.

Most *Cinara* species host only two obligate partners. However, in some species, *Type-X* was found with both *Serratia* and *Buchnera* in all the individuals sampled (Fig. 1). This either suggests that some *Cinara* species have more than two obligate symbionts, as observed in other arthropods (Koga *et al.* 2013), or that these species are currently in a transitional state, in which the ancestral *Serratia* (Fig. S9) has not yet been eliminated. A third possibility is that one of the co-symbionts represents a more recent facultative infection. This may apply to the species hosting *Wolbachia.* Although this bacterium is usually found in all individuals of the species it infects (Fig. S4) and associated with *Serratia, Wolbachia* has not established a long-term association with *Cinara*, as it is not fixed in any particular clade of the genus. *Wolbachia* may probably be a facultative symbiont that is widespread in aphids’ populations thanks to its ability to manipulate reproduction, as previously suggested (Augustinos *et al.* 2011; Gomez-Valero *et al.* 2004). We have detected many individuals in which the cosymbionts described above are present together with other bacteria (*Arsenophonus, Edwardsiella, Regiella, Rickettsia, Hamiltonella* or *Spiroplasma;* Fig. 1, S3). These additional partners are probably facultative infections, because they occur sporadically within the species and their association with *Cinara* is not evolutionarily stable (Fig. S4, Table S4).

### Evolutionary history of symbiotic associations

Two evolutionary scenarios could explain the distribution of co-symbionts observed here. *Serratia* may have infected an ancestor of the Lachninae (Fig. S9), a subfamily that originated more than 70 million years ago (Chen *et al.* 2015; Meseguer *et al.* 2015). The division of labor in the nutrition of *Cinara* may have been established then (Fig. 2) (Lamelas *et al.* 2011a; Lamelas *et al.* 2011b). At various times points between the Oligocene and the present, *Serratia* may then have been lost from several clades of *Cinara* (Fig. S9). These losses were associated with the acquisition of an alternative co-resident, *Erwinia, Type-X* or *Sodalis.* This scenario implies a long history of cospeciation between *Cinara* and *Serratia*, followed by further cospeciation between *Cinara* and its more recently acquired cosymbionts. Alternatively, there may have been multiple independent colonizations of an already diversified *Cinara* genus by several lineages of *Serratia, Erwinia, Type-X* and *Sodalis.*

We cannot conduct robust cospeciation tests using endosymbiont phylogenies based on the short 16S rRNA marker sequenced here. However, the patterns of specificity revealed here shed some light on the history of the symbiotic associations between *Cinara* and its bacterial symbionts. The distribution of 16S *Serratia* haplotypes across *Cinara* species suggests that the codiversification history of *Serratia* and *Cinara* probably involved both cospeciation and multiple infections. On the one hand, the interaction between the two partners is generally species-specific (Fig. S5, Table 1), which is consistent with cospeciation scenarios. On the other hand, several *Serratia* strains are common to distantly related aphid species (e.g. one *Serratia* haplotype is present in 34 unrelated *Cinara* species), and most of the species containing these haplotypes are also infected with another *Serratia* strain. Furthermore, previous studies suggest that the *Serratia* strains associated with *C. cedri* and *C. tujafilina* could belong to two distantly related lineages (Manzano-Marin & Latorre 2014; Manzano-Marin *et al.* 2016a). Altogether, these findings demonstrate that, during the course of its evolution, *Cinara* has experienced multiple infections with different lineages of *Serratia.* Some lineages may have infected the ancestors of *Cinara*, establishing an obligate association and partly cospeciated with them, whereas other *Serratia* lineages may have infected different species of the genus more recently. Thus obligate and facultative *Serratia* strains might even be found co-infecting the same aphid (Fig. S5). This would explain why previous studies based on phylogenetic analyses of *Serratia* 16S rDNA sequences rejected the hypothesis of cospeciation between *Serratia* and *Cinara* (Burke *et al.* 2009; Lamelas *et al.* 2008).

Similarly, the overall patterns of *Type-X* and *Sodalis* specificity for their hosts suggest that these associations may result from a combination of cospeciation and host-switches (Fig. S5). Codiversification scenarios will be difficult to unravel for these associations, as the presence of multiple, slightly divergent, bacterial haplotypes in each aphid specimen (Appendix 2) suggests some co-infections by several strains, and/or that the 16S rRNA gene is present in multiple copies in a single symbiont type (Koga *et al.* 2013). The association of *Cinara* and *Erwinia* is not species-specific, although each *Cinara* species generally contains only one *Erwinia* haplotype, closely related aphid species harbor *Erwinia* strains of the same haplotype. This suggests a lack of differentiation in *Erwinia* during the speciation of aphids, or that the DNA fragment studied here may not have been variable enough to reflect interspecific variation.

The patterns of symbiont-aphid specificity depicted here show that future studies should make use of several single-copy bacterial DNA markers and take into account coinfections by several strains and possibly bacterial cell polyploidy-*i.e.* the presence of many, slightly divergent, bacterial chromosomes in a single bacteriocyte (Komaki & Ishikawa 1999)-in the investigation of codiversification history between *Cinara* and its obligate symbionts.

### Which factors drive symbiont replacement?

The repeated evolution of dual symbiosis in *Cinara* provides us with a unique opportunity to investigate the factors favoring one co-resident symbiont over another in symbiotic associations between aphids and bacteria. The presence of facultative symbionts in aphid populations, and insects in general, is generally explained by the ecological niche occupied by their hosts (Henry *et al.* 2015; Liberti *et al.* 2015; Oliver *et al.* 2010). However, little is currently known about the forces governing the evolution of these associations when the symbionts are essential for the system to survive. We found that changes in obligate co-symbionts were not correlated with evolutionary transitions in *Cinara* (Table S5). All of the obligate co-symbionts identified here were found in aphid species using a wide range of ecological niches (Fig. 3, 4). This implies that the acquisition of new cosymbionts did not trigger the adaptation of the host aphid to environmental conditions. The addition of a new partner to symbiotic associations could be explained by a degradation of the functions of the existing partner (e.g. due to genetic drift) (Bennett & Moran 2015). In a dual symbiotic systems in leafhoppers (Bennett *et al.* 2014), the most recently recruited symbiont has been shown to possess a less stable genome than the ancient symbiont and has been repeatedly replaced. A similar situation may apply to our model system; a loss of symbiotic functions in *Serratia* may have favored the establishment of an alternative bacterium, while *Buchnera* persisted in the association. Alternatively, the relatively rapid turnover of cosymbionts alongside *Buchnera* may result from competitive interactions between bacteria with similar metabolic capabilities. Essential functions have been lost from *Buchnera* in an ancestor of Lachninae (Fig. 2, S9) (Manzano-Marin *et al.* 2016a). The delegation of these symbiotic functions of *Buchnera* to a bacterial partner may have paved the way for colonization by any bacterium with similar metabolic capacities. In this scenario, the identity of the new partner depends on the outcome of competition between bacteria, regulated by their population dynamics (*i.e*., demographic advantages due to higher replication rates) rather than the selective advantages they confer on the host.

**Figure 4.**
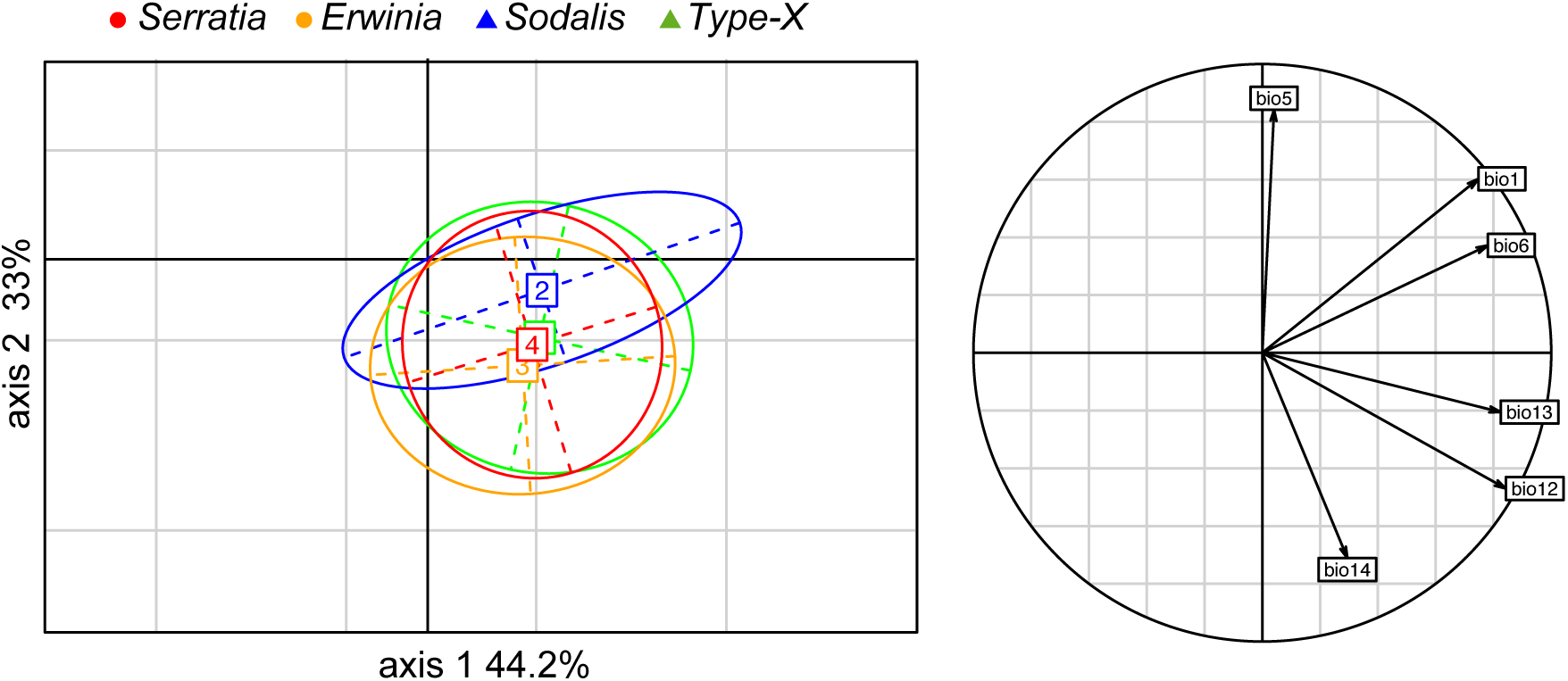
Circle of correlation and factor scores obtained from a principal component analysis performed on bioclimatic variables. Ellipses delimit the climatic envelopes of aphids hosting the different cosymbionts. Climatic values were extracted from the localities where aphid specimens carrying the essential symbionts were found. The percentages of variance explained by the first two principal components are indicated in the axis labels. Abbreviations; BIO1 = Annual Mean Temperature; BIO5 = Max Temperature of Warmest Month; BIO6 = Min Temperature of Coldest Month; BIO12 = Annual Precipitation; BIO13 = Precipitation of Wettest Month; BIO14 = Precipitation of Driest Month

## Conclusions

We report here a highly dynamic di-symbiotic system in the second most diverse aphid genus. An additional endosymbiotic partner is required in *Cinara* to complement *Buchnera* in its nutritional role. Interestingly, this new “flatmate” has been repeatedly replaced during the diversification of the group but it never replaces *Buchnera.* This mirrors findings for other sap-feeding insects where the primary symbiont is supplemented by an additional bacterium. In Auchenorrhyncha, the ancestral symbiont, *Sulcia*, coexists with another obligate symbiont, the identity of which differs between lineages (Koga *et al.* 2013). In some mealybugs, the primary symbiont shelter within its cells another Gammaproteobacterium symbiont that has also been repeatedly replaced (Husnik & McCutcheon 2016). The aphid sister group (Adelgidae), has not established a long-term association with a primary symbiont, but is found associated with a diverse set of obligate bacterial partners throughout its evolutionary history (Toenshoff *et al.* 2012; Toenshoff *et al.* 2014). Our results suggest that the succession of essential symbionts does not necessarily result from the adaptation of their hosts to changing ecological conditions. It might therefore be driven by factors intrinsic to the evolutionary dynamics of endosymbionts, such as rapid genome deterioration or the competitive displacement of symbionts providing similar benefits to the host.

## Authors Contributions

EJ, ASM, ACA designed the study. ASM and EJ wrote the paper with contributions of AMM. ASM, AMM and MG analysed the data. ALC performed the molecular work. ACA identified the aphid specimens. ACA and EJ collected the specimens.

## Acknowledgements

Research funding was provided by a Marie-Curie FP7-COFUND (AgreenSkills fellowship-26719) to ASM, the ANR Phylospace and the project *(“Cinara’s* microbiome”) from the Agropolis foundation/Labex Agro to EJ. The authors are grateful to the CBGP-HPC computational platform, J. Abbat and S. Joly for help with statistical analyses. To F. Condamine, F. Kjellberg and the three anonymous reviewers for comments on the manuscript.

## Data Accessibility

The 16S rRNA sequence data produced in this study are available from the Dryad Digital Repository (doi:10.5061/dryad.9jn6r). Appendices 1-3 are also accessible in Dryad (doi:10.5061/dryad.sk130); Appendix 1 includes collection details for aphid samples and GenBank accession numbers. Appendix 2 shows a contingency table reporting bacterial sequence occurrences across aphid samples, and Appendix 3 includes a summary of the number of reads and unique sequences per sample. DNA sequences for the aphid phylogeny are available in Genbank accessions KY064183-KY064504. Reads used for genome assembly as well as draft assemblies for the endosymbionts’ genomes have been deposited in the European Nucleotide Archive under the project numbers PRJEB15400 (*C. fornacula*), PRJEB15504 (*C. confinis)*, PRJEB15506 (*C. pseudotaxifoliae*) and PRJEB15507 (*C. strobi).*

## References

Augustinos AA, Santos-Garcia D, Dionyssopoulou E, et al. (2011) Detection and characterization of *Wolbachia* infections in natural populations of Aphids: Is the hidden diversity fully unraveled? Plos One 6, e28695.

Bankevich A, Nurk S, Antipov D, et al. (2012) SPAdes: a new genome assembly algorithm and its applications to single-cell sequencing. Journal of Computational Biology 19, 455–477.

Bennett GM, McCutcheon JP, MacDonald BR, Romanovicz D, Moran NA (2014) Differential genome evolution between companion symbionts in an insect-bacterial symbiosis. Mbio 5.

Bennett GM, Moran NA (2015) Heritable symbiosis: The advantages and perils of an evolutionary rabbit hole. Proceedings of the National Academy of Sciences 112, 10169–10176.

Blackman RL, Eastop VF (1994) Aphids on the world trees: an identification and information guide The Natural History Museum, London, UK.

Blackman RL, Eastop VF (2000) Aphids on the world’s crops: an identification and information guide, Second edition edn. Willey and sons, New York.

Brown JM, Hedtke SM, Lemmon AR, Lemmon EM (2010) When trees grow too long: Investigating the causes of highly inaccurate bayesian branch-length estimates. Systematic Biology QP, 145–161.

Buchner P (1965) Endosymbiosis of animal with plant microorganims Interscience, New York.

Burke GR, Normark BB, Favret C, Moran NA (2009) Evolution and diversity of facultative symbionts from the aphid subfamily Lachninae. Applied and Environmental Microbiology SQ, 5328–5335.

Caporaso JG, Kuczynski J, Stombaugh J, et al. (2010) QIIME allows analysis of high-throughput community sequencing data. Nature Methods S, 335–336.

Cayetano L, Vorburger C (2015) Symbiont-conferred protection against Hymenopteran parasitoids in aphids: how general is it? Ecological Entomology TU, 85–93.

Charles H, Ishikawa A (1999) Physical and genetic map of the genome of Buchnera, the primary endosymbiont of the pea aphid Acyrthosiphon pisum. Journal of Molecular Evolution 48, 142–150.

Chen R, Favret C, Jiang L, Wang Z, Qiao G (2015) An aphid lineage maintains a bark-feeding niche while switching to and diversifying on conifers. Cladistics MWI9 KU=KKKKXcla=KRK4K.

Clark EL, Daniell TJ, Wishart J, Hubbard SF, Karley AJ (2012) How conserved are the bacterial communities associated with aphids? A detailed assessment of the *Brevicoryne brassicae*(Hemiptera: Aphididae) using 16S rDNA. Environmental Entomology 4K, 1386–1397.

Coeur d’acier A, Cruaud A, Artige E, et al. (2014) DNA barcoding and the associated PhylAphidB@se website for the identification of European aphids (Insecta: Hemiptera: Aphididae). Plos One 9, e97620.

Conord C, Despres L, Vallier A, et al. (2008) Long-term evolutionary stability of bacterial endosymbiosis in curculionoidea: Additional evidence of symbiont replacement in the dryophthoridae family. Molecular Biology and Evolution R5, 859–868.

Dolédec S, Chessel D (1987) Rythmes saisonniers et composantes stationnelles en milieu aquatique. I: Description d’un plan d’observation complet par projection de variables. Acta Oecologia 8, 403–426.

Dormann CF, Gruber B, Fruend J (2008) Introducing the bipartite package: analysing ecological networks. R news 8, 8–11.

Douglas AE (1998) Nutritional interactions in insect-microbial symbioses: aphids and their symbiotic bacteria Buchnera. Annual Review of Entomology 43, 17–37.

Dray S, Dufour A-B (2007) The ade4 package: implementing the duality diagram for ecologists. Journal of Statistical Software RR, 1–20.

Drummond AJ, Ashton B, Buxton S, et al. (2010) Geneious v5.5. Available from http://www.geneious.com. Biomatters, Auckland, New Zealand.

Edgar RC, Haas BJ, Clemente JC, Quince C, Knight R (2011) UCHIME improves sensitivity and speed of chimera detection. Bioinformatics R7, 2194–2200.

Favret C, Voegtlin DJ (2004) A revision of the *Cinara* species (Hemiptera: Aphididae) of the United States Pinyon pines. Annals of the Entomological Society of America 97, 1165–1197.

Flandrois J-P, Perrière G, Gouy M (2014) leBIBIQBPP: A set of databases and a webtool for automatic phylogenetic analysis of prokaryotic sequences. http://umr5558-bibiserv.univ-lyon1.fr/lebibi/lebibi.cgi

Frago E, Dicke M, Godfray HCJ (2012) Insect symbionts as hidden players in insect-plant interactions. Trends in Ecology & Evolution R7, 705–711.

Goecks J, Nekrutenko A, Taylor J, Team TG (2010) Galaxy: a comprehensive approach for supporting accessible, reproducible, and transparent computational research in the life sciences. Genome Biology KK, R86.

Gomez-Valero L, Soriano-Navarro M, Perez-Brocal V, et al. (2004) Coexistence of *Wolbachia* with *Buchnera aphidicola* and a secondary symbiont in the aphid *Cinara cedri*. Journal of Bacteriology K86, 6626–6633.

Gosalbes MJ, Lamelas A, Moya A, Latorre A (2008) The striking case of tryptophan provision in the cedar aphid Cinara cedri. Journal of Bacteriology K9U, 6026–6029.

Guay JF, Boudreault S, Michaud D, Cloutier C (2009) Impact of environmental stress on aphid clonal resistance to parasitoids: Role of *Hamiltonella defensa* bacterial symbiosis in association with a new facultative symbiont of the pea aphid. Journal of Insect Physiololy 55, 919–926.

Hansen AK, Moran NA (2014) The impact of microbial symbionts on host plant utilization by herbivorous insects. Molecular Ecology R3, 1473–1496.

Harada H, Oyaizu H, Kosako Y, Ishikawa H (1997) *Erwinia aphidicola*, a new species isolated from pea aphid, Acyrthosiphon pisum. Journal of General and Applied Microbiology 43, 349–354.

Harmon LJ, Weir JT, Brock CD, Glor RE, Challenger W (2008) GEIGER: investigating evolutionary radiations. Bioinformatics R4, 129–131.

Henry LM, Maiden MCJ, Ferrari J, Godfray HCJ (2015) Insect life history and the evolution of bacterial mutualism. Ecology Letters 18, 516–525.

Hijmans RJ, Cameron SE, Parra JL, Jones PG, Jarvis A (2005) Very high resolution interpolated climate surfaces for global land areas. International journal of Climatology 25, 1965–1978.

Ho LST, Ané C (2014) A linear-time algorithm for Gaussian and non-Gaussian trait evolution models. Systematic Biology 63, 397–408.

Huelsenbeck JP, Larget B, Alfaro ME (2004) Bayesian phylogenetic model selection using reversible jump Markov chain Monte Carlo. Molecular Biology and Evolution 21, 1123–1133.

Husnik F, McCutcheon JP (2016) Repeated replacement of an intrabacterial symbiont in the tripartite nested mealybug symbiosis. Proceedings of the National Academy of Sciences 113, E5416–E5424.

Ives AR, Garland T (2010) Phylogenetic logistic regression for binary dependent variables. Systematic Biology 59, 9–26.

Jousselin E, Clamens AL, Galan M, et al. (2016) Assessment of a 16S rRNA amplicon Illumina sequencing procedure for studying the microbiome of a symbiont-rich aphid genus. Molecular Ecology Resources, n/a-n/a.

Jousselin E, Cruaud A, Genson G, et al. (2013) Is ecological speciation a major trend in aphids? Insights from a molecular phylogeny of the conifer-feeding genus *Cinara*. Frontiers in Zoology 10, 56–73.

Katoh K, Toh H (2008) Recent developments in the MAFFT multiple sequence alignment program. Briefings in Bioinformatics 9, 286–298.

Koga R, Bennett GM, Cryan JR, Moran NA (2013) Evolutionary replacement of obligate symbionts in an ancient and diverse insect lineage. Environmental microbiology 15, 2073–2081.

Koga R, Moran NA (2014) Swapping symbionts in spittlebugs: evolutionary replacement of a reduced genome symbiont. Isme Journal 8, 1237–1246.

Komaki K, Ishikawa H (1999) Intracellular bacterial symbionts of aphids possess many genomic copies per bacterium. Journal of Molecular Evolution 48, 717–722.

Kozich JJ, Westcott SL, Baxter NT, Highlander SK, Schloss PD (2013) Development of a dual-index sequencing strategy and curation pipeline for analyzing amplicon sequence data on the MiSeq Illumina sequencing platform. Applied and Environmental Microbiology 79, 5112–5120.

Lamelas A, Gosalbes MJ, Manzano-Marin A, et al. (2011a) *Serratia symbiotica* from the aphid *Cinara cedri:* a missing link from facultative to obligate insect endosymbiont. Plos Genetics 7.

Lamelas A, Gosalbes MJ, Moya A, Latorre A (2011b) New clues about the evolutionary history of metabolic losses in bacterial endosymbionts, provided by the genome of *Buchnera aphidicola* from the aphid *Cinara tujafilina*. Applied and Environmental Microbiology 77, 4446–4454.

Lamelas A, Perez-Brocal V, Gomez-Valero L, et al. (2008) Evolution of the secondary symbiont *“Candidatus Serratia symbiotica”* in aphid species of the subfamily Lachninae. Applied and Environmental Microbiology 74, 4236–4240.

Lanfear R, Calcott B, Ho SYW, Guindon S (2012) PartitionFinder: combined selection of partitioning schemes and substitution models for phylogenetic analyses. Molecular Biology and Evolution 29, 1695–1701.

Langmead B, Salzberg SL (2012) Fast gapped-read alignment with Bowtie 2. Nature Methods 9, 357–U354.

Liberti J, Sapountzis P, Hansen LH, et al. (2015) Bacterial symbiont sharing in Megalomyrmex social parasites and their fungus-growing ant hosts. Molecular Ecology 24, 3151–3169.

Manzano-Marin A, Latorre A (2014) Settling down: the genome of *Serratia symbiotica* from the aphid *Cinara tujafilina* zooms in on the process of accommodation to a cooperative intracellular life. Genome Biol Evol 6, 1683–1698.

Manzano-Marin A, Simon J-C, Latorre A (2016a) Reinventing the Wheel and Making It Round Again: Evolutionary Convergence in Buchnera - Serratia Symbiotic Consortia between the Distantly Related Lachninae Aphids Tuberolachnus salignus and Cinara cedri. Genome Biology and Evolution V, 1440–1458.

Manzano-Marin A, Szabo G, Simon J-C, Horn M, Latorre A (2016b) Happens in the best of subfamilies: Replacement and internalisation of co-obligate *Serratia* endosymbionts in Lachninae aphids. bioRxivorg, 059816.

McCutcheon JP, Moran NA (2010) Functional convergence in reduced genomes of bacterial symbionts spanning 200 My of evolution. Genome Biology and Evolution R, 708–718.

McCutcheon JP, Moran NA (2012) Extreme genome reduction in symbiotic bacteria. Nature Reviews Microbiology KU, 13–26.

McCutcheon John P, von Dohlen Carol D An interdependent metabolic patchwork in the nested symbiosis of Mealybugs. Current Biology RK, 1366–1372.

Meseguer AS, Aldasoro JJ, Sanmartín I (2013) Bayesian inference of phylogeny, morphology and range evolution reveals a complex evolutionary history in St. John’s wort *(Hypericum)*. Molecular Phylogenetics and Evolution 6S, 379–403.

Meseguer AS, Coeur d’acier A, Genson G, Jousselin E (2015) Unravelling the historical biogeography and diversification dynamics of a highly diverse conifer-feeding aphid genus. Journal of Biogeography TR, 1482–1492.

Mizrahi-Man O, Davenport ER, Gilad Y (2013) Taxonomic Classification of Bacterial 16S rRNA Genes Using Short Sequencing Reads: Evaluation of Effective Study Designs. Plos One V.

Moran NA (1996) Accelerated evolution and Muller’s rachet in endosymbiotic bacteria. Proceedings of the National Academy of Sciences of the United States of America PY, 2873–2878.

Moran NA, Telang A (1998) Bacteriocyte-associated symbionts of insects - A variety of insect groups harbor ancient prokaryotic endosymbionts. Bioscience TV, 295–304.

Oakeson KF, Gil R, Clayton AL, et al. (2014) Genome degeneration and adaptation in a nascent stage of symbiosis. Genome Biology and Evolution L, 76–93.

Oliver KM, Campos J, Moran NA, Hunter MS (2008) Population dynamics of defensive symbionts in aphids. Proceedings of the Royal Society B-Biological Sciences RSQ, 293–299.

Oliver KM, Degnan PH, Burke GR, Moran NA (2010) Facultative symbionts in aphids and the horizontal transfer of ecologically important traits. Annual Review of Entomology QQ, 247–266.

Oliver KM, Martinez AJ (2014) How resident microbes modulate ecologically-important traits of insects. Current Opinion in Insect Science T, 1–7.

Pagel M (1994) Detecting correlated evolution on phylogenies: a general method for the comparative analysis of discrete characters. Proceedings of the Royal Society B-Biological Sciences RQQ, 3745.

Paradis E, Claude J, Strimmer K (2004) APE: Analyses of phylogenetics and evolution in R language. Bioinformatics RU, 289–290.

Perez-Brocal V, Gil R, Ramos S, et al. (2006) A small microbial genome: The end of a long symbiotic relationship? Science YKT, 312–313.

Quast C, Pruesse E, Yilmaz P, et al. (2013) The SILVA ribosomal RNA gene database project: improved data processing and web-based tools. Nucleic Acids Research TK, D590–D596.

Rambaut A (2002) Se-Al: sequence alignment editor. Available: http://tree.bio.ed.ac.uk/software/seal/.

Rispe C, Moran NA (2000) Accumulation of deleterious mutations in endosymbionts: Muller’s ratchet with two levels of selection. American Naturalist WL, 425–441.

Ronquist F, Teslenko M, Van Der Mark P, et al. (2012) Mrbayes 3.2: Efficient bayesian phylogenetic inference and model choice across a large model space. Systematic Biology LK, 539–542.

Russell JA, Moran NA (2006) Costs and benefits of symbiont infection in aphids: variation among symbionts and across temperatures. Proceedings of the Royal Society B-Biological Sciences RSY, 603–610.

Schloss PD, Westcott SL (2011) Assessing and improving methods used in operational taxonomic unit-based approaches for 16S rRNA gene sequence analysis. Applied and Environmental Microbiology 77, 3219–3226.

Schmieder R, Edwards R (2011) Quality control and preprocessing of metagenomic datasets. Bioinformatics 27, 863–864.

Simonet P, Duport G, Gaget K, et al. (2016) Direct fow cytometry measurements reveal a fine-tuning of symbiotic cell dynamics according to the host developmental needs in aphid symbiosis. Scientific Reports 6, 19967.

Smith AH, tukasik P, O’Connor MP, et al. (2015) Patterns, causes and consequences of defensive microbiome dynamics across multiple scales. Molecular Ecology 24, 1135–1149.

Smith W, Oakeson K, Johnson K, et al. (2013) Phylogenetic analysis of symbionts in feather-feeding lice of the genus Columbicola: evidence for repeated symbiont replacements. BMC Evolutionary Biology 13, 109.

Snyder AK, McMillen CM, Wallenhorst P, Rio RVM (2011) The phylogeny of *Sodalis-like* symbionts as reconstructed using surface-encoding loci. FEMS Microbiology Letters 317, 143–151.

Takiya DM, Tran PL, Dietrich CH, Moran NA (2006) Co-cladogenesis spanning three phyla: leafhoppers (Insecta: Hemiptera: Cicadellidae) and their dual bacterial symbionts. Molecular Ecology 15, 4175–4191.

Toenshoff ER, Gruber D, Horn M (2012) Co-evolution and symbiont replacement shaped the symbiosis between adelgids (Hemiptera: Adelgidae) and their bacterial symbionts. Environmental microbiology 14, 1284–1295.

Toenshoff ER, Szabo G, Gruber D, Horn M (2014) The pine bark adelgid, *Pineus strobi*, contains two novel bacteriocyte-associated Gammaproteobacterial symbionts. Applied and Environmental Microbiology 80, 878–885.

Toft C, Andersson SGE (2010) Evolutionary microbial genomics: insights into bacterial host adaptation. Nature Reviews Genetics 11, 465–475.

Toju H, Tanabe AS, Notsu Y, Sota T, Fukatsu T (2013) Diversification of endosymbiosis: replacements, co-speciation and promiscuity of bacteriocyte symbionts in weevils. Isme Journal 7, 1378–1390.

Vigneron A, Masson F, Vallier A, et al. (2014) Insects recycle endosymbionts when the benefit Is over. Current Biology 24, 2267–2273.

Warnes GR, Bolker B, Bonebakker L, et al. (2015) gplots: various R programming tools for plotting data. R package.

Wernegreen JJ (2012) Mutualism meltdown in insects: bacteria constrain thermal adaptation. Current Opinion in Microbiology 15, 255–262.

Wilson ACC, Ashton PD, Calevro F, et al. (2010) Genomic insight into the amino acid relations of the pea aphid, *Acyrthosiphon pisum*, with its symbiotic bacterium *Buchnera aphidicola*. Insect Molecular Biology 19, 249–258.

Wu D, Daugherty SC, Van Aken SE, et al. (2006) Metabolic complementarity and genomics of the dual bacterial symbiosis of sharpshooters. PLoS Biology 4, 1079–1092.

Zachos JC, Dickens GR, Zeebe RE (2008) An early Cenozoic perspective on greenhouse warming and carbon-cycle dynamics. Nature 451, 279–283.

Zhang J, Kapli P, Pavlidis P, Stamatakis A (2013) A general species delimitation method with applications to phylogenetic placements. Bioinformatics 29, 2869–2876.

Zytynska SE, Weisser WW (2016) The natural occurrence of secondary bacterial symbionts in aphids. Ecological Entomology 41, 13–26.

